## Supplemental Figures and Note for "Frequency-dependent selection of neoantigens fosters tumor immune escape and predicts immunotherapy response"

### 1 Supplementary Figures

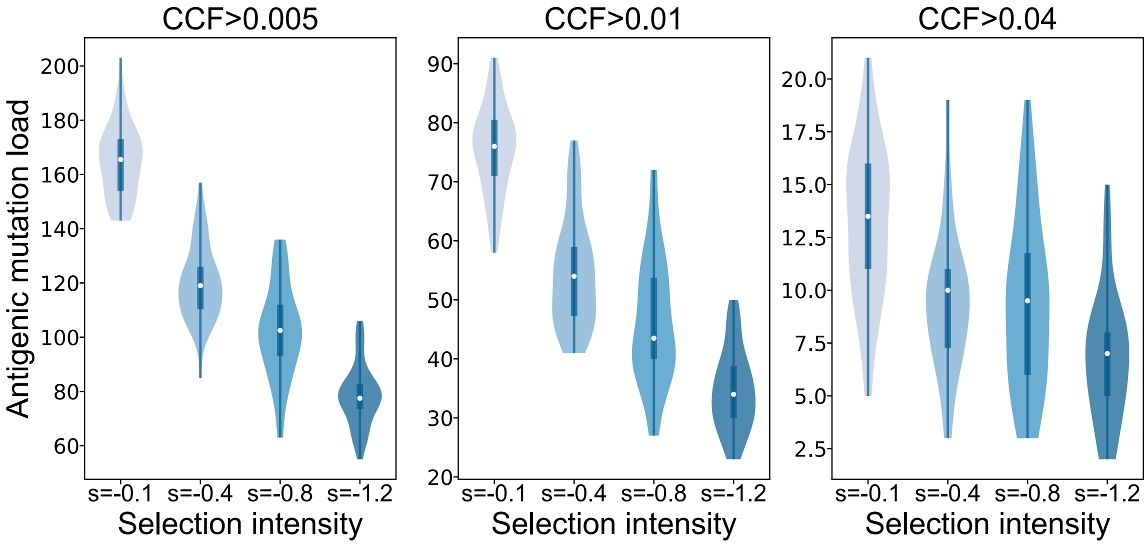


**Supplementary Fig. 1. Neoantigen loads with different CCF cutoffs.** Number of neoantigens accumulated in simulated tumors undergoing PS at varying selective intensities (each with 50 simulations). Cutoffs of neoantigen clonality CCF$>$0.005, 0.01 and 0.04 are used respectively.

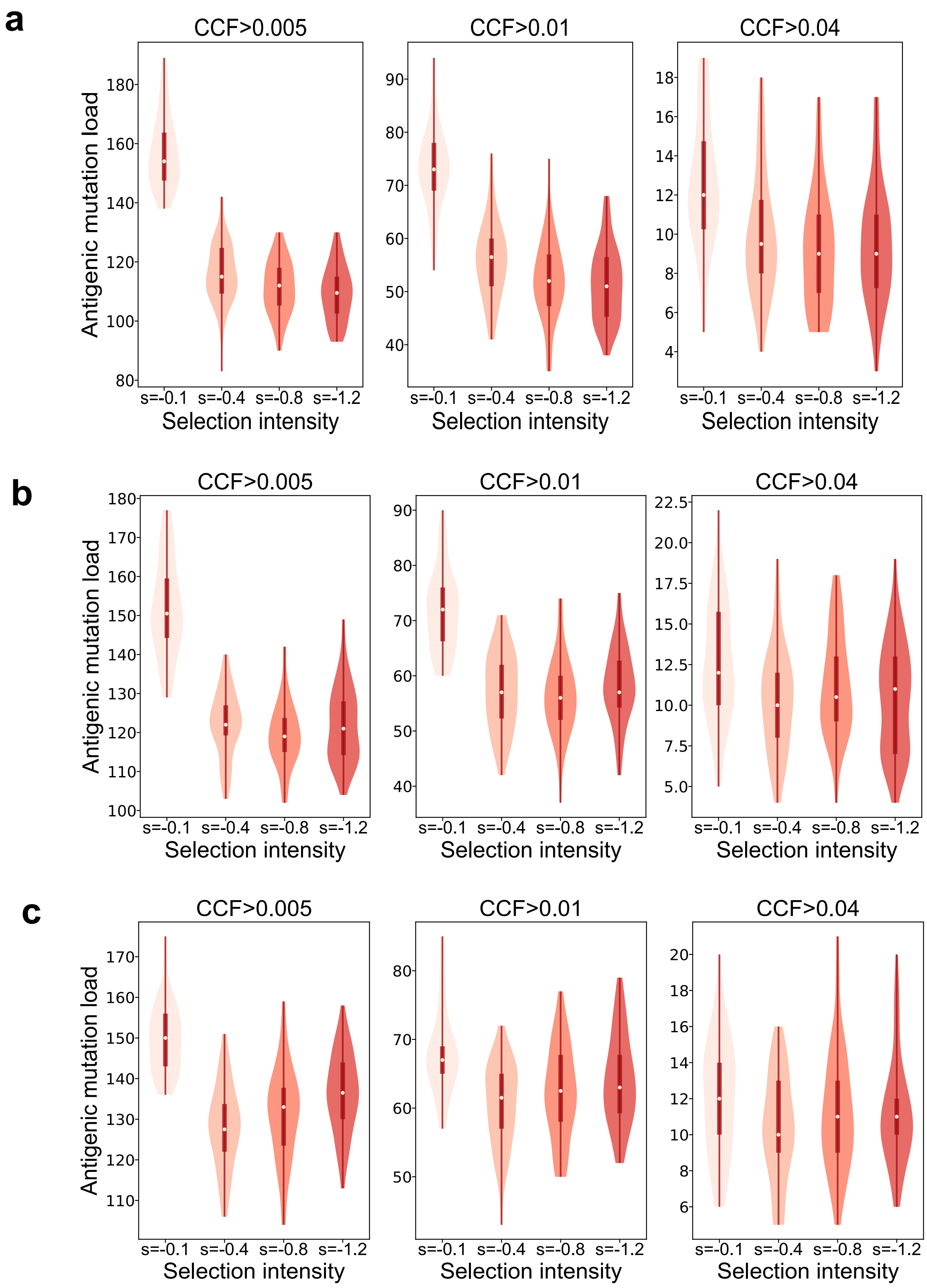


**Supplementary Fig. 2. Neoantigen loads with different cell immunogenicity cutoffs. a-c,** Number of neoantigens accumulated in simulated tumors at varying selective intensities (each with 50 simulations). Simulated tumors undergoing NFDS with cell immunogenicity threshold ($c_1$) equals 0.3 (**a**), 0.5 (**b**) and 0.7 (**c**) are shown. Neoantigens with CCF$>$0.005,0.01,0.04 are used, respectively.

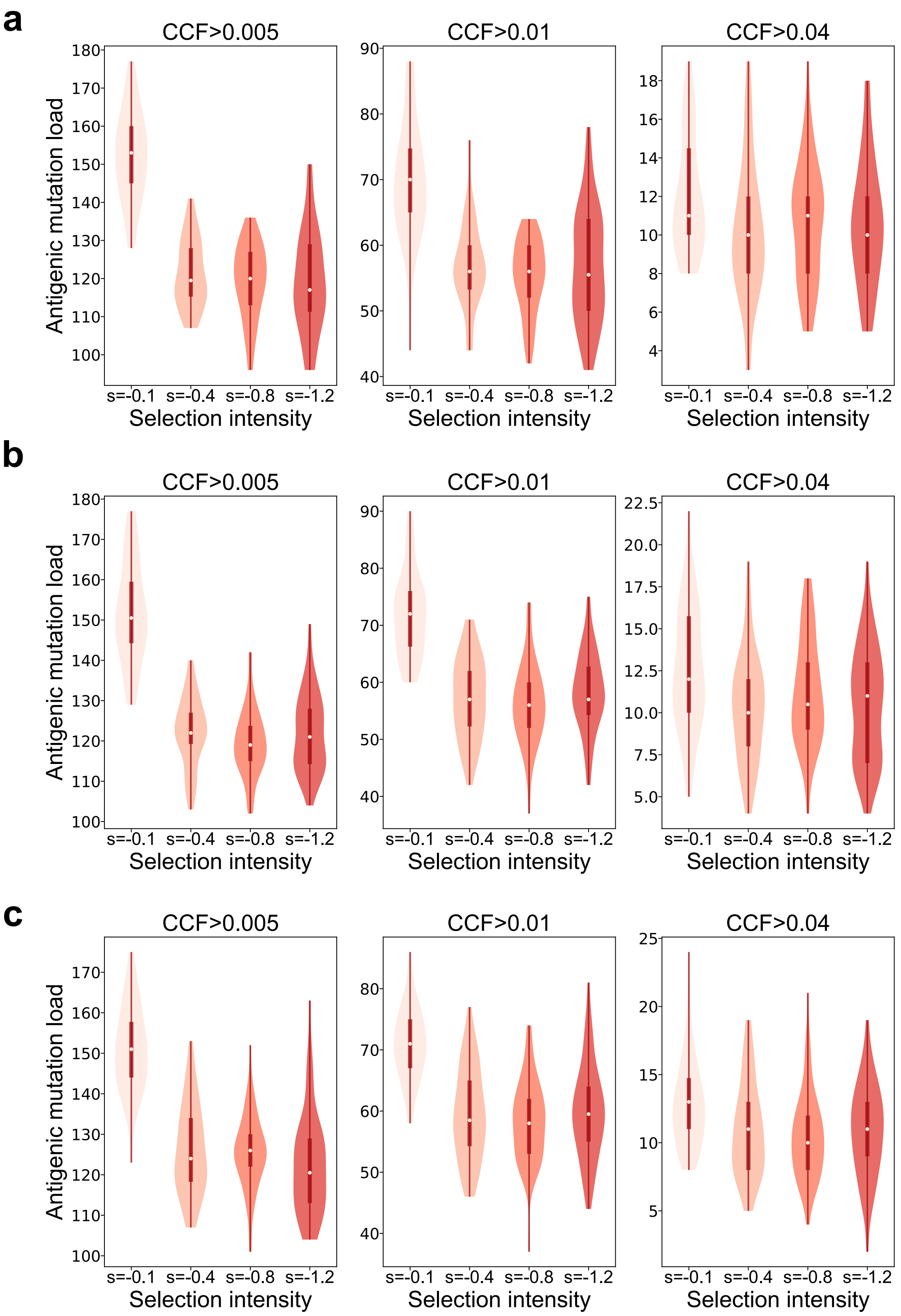


**Supplementary Fig. 3. Neoantigen loads with different tumor immunogenicity cutoffs. a-c,** Number of neoantigens accumulated in simulated tumors at varying selective intensities (each with 50 simulations). Simulated tumors undergoing NFDS with cell immunogenicity threshold ($c_2$) equals 0.3 (**a**), 0.5 (**b**) and 0.7 (**c**) are shown. Neoantigens with CCF$>$0.005,0.01,0.04 are used, respectively.

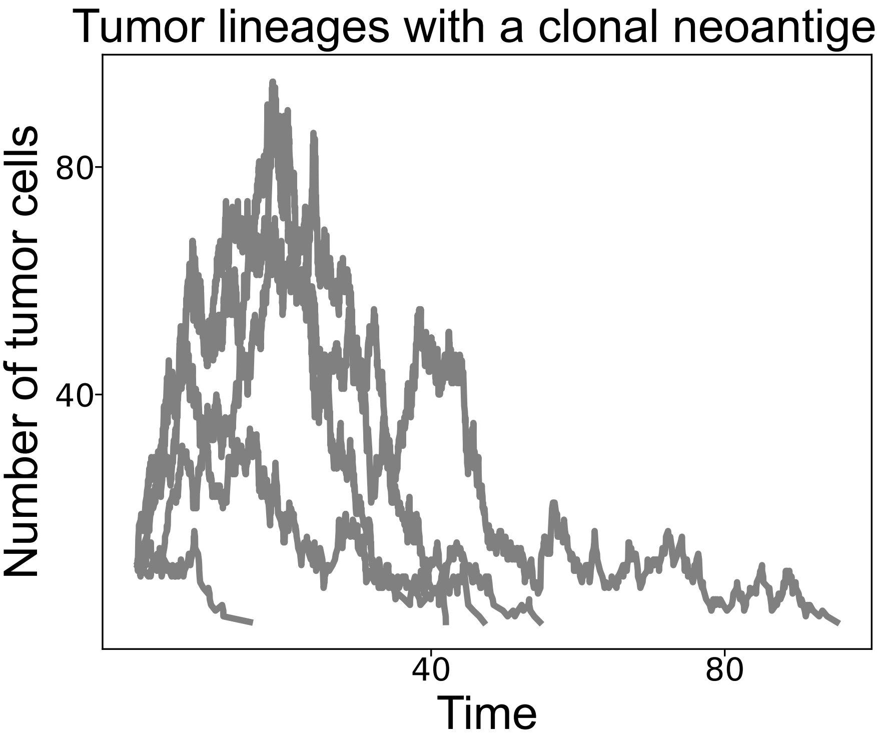


**Supplementary Fig. 4. Tumor lineages with a clonal neoantigen.** Five simulated tumors that each contains one clonal antigenic mutation with antigenicity score equals to 0.2 and mutation rate of $\mu = 5$ at selective intensity $s = -0.8$.

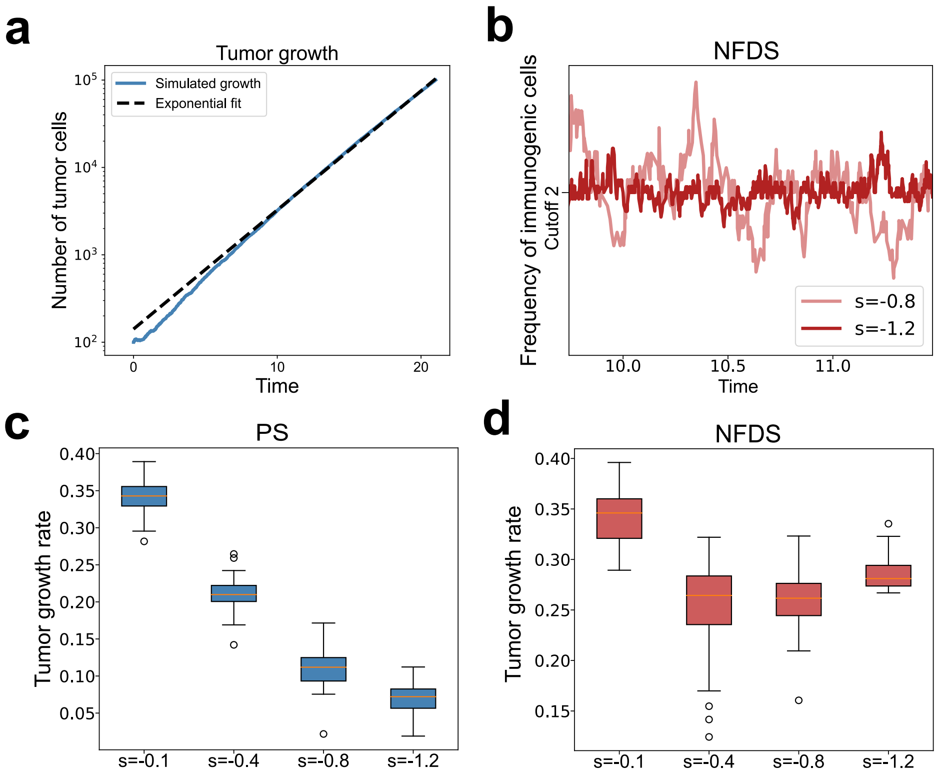


**Supplementary Fig. 5. Tumor growth rates under PS and NFDS, respectively. a,** Exponential fitting of simulated tumor growth. **b,** Frequency curves of immunogenic cells of two simulated tumors under negative frequency-dependent selection (NFDS) with selective intensity $s = 0.8$ and 1.2, respectively. **c-d,** Tumor growth rate at varying selective intensities in simulated tumors ($n = 50$) undergoing PS and NFDS, respectively.

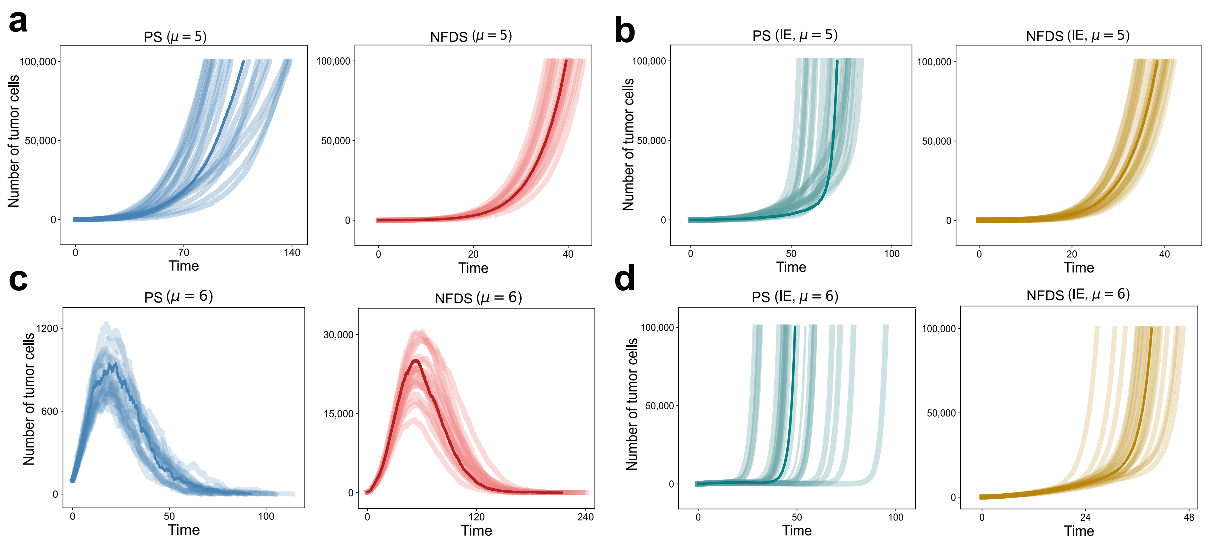


**Supplementary Fig. 6. Tumor growth curves with different mutation rates.** **a,** Growth curves of 20 simulated tumors under PS and NFDS, respectively. **b,** Growth curves of 20 simulated tumors under PS (subclonal immune escape or IE) and NFDS (subclonal IE), respectively. Here mutation rate $\mu = 5$ and selection coefficient $s = -0.8$ are used. **c,** Growth curves of 20 simulated tumors under PS and NFDS, respectively. **d,** Growth curves of 20 simulated tumors under PS (subclonal immune escape or IE) and NFDS (subclonal IE), respectively. Here mutation rate $\mu = 6$ and selection coefficient $s = -0.8$ are used.

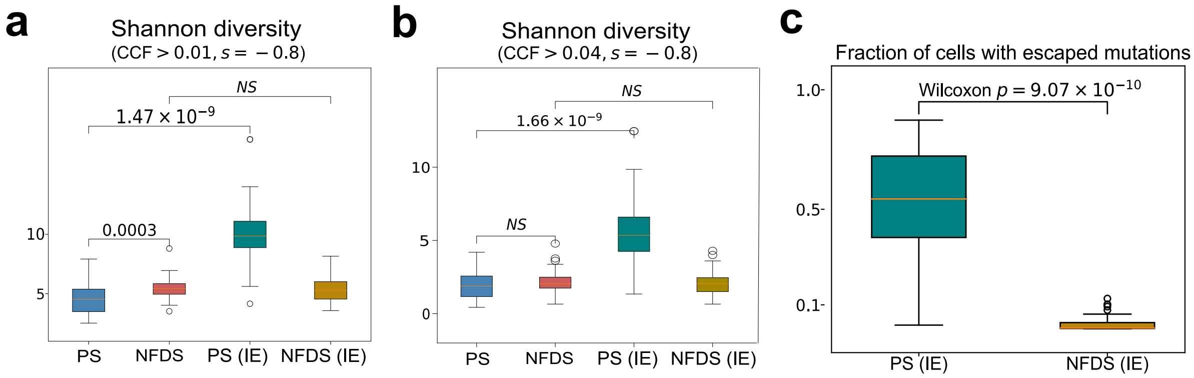


**Supplementary Fig. 7. Shannon diversity and fraction of immune-escaped cells a-b,** Box plots showing the antigenic mutations accumulated in 50 simulated tumors with cancer cell fraction (CCF)$> 0.01$ (**a**) and CCF$> 0.04$ (**b**), respectively. The Shannon diversity of antigenic mutations from 50 simulated tumors under four evolutionary scenarios are shown. **c,** Box plots of fractions of cells with escaped mutations in 50 simulated tumors under PS and NFDS with subclonal immune escape (IE), respectively.

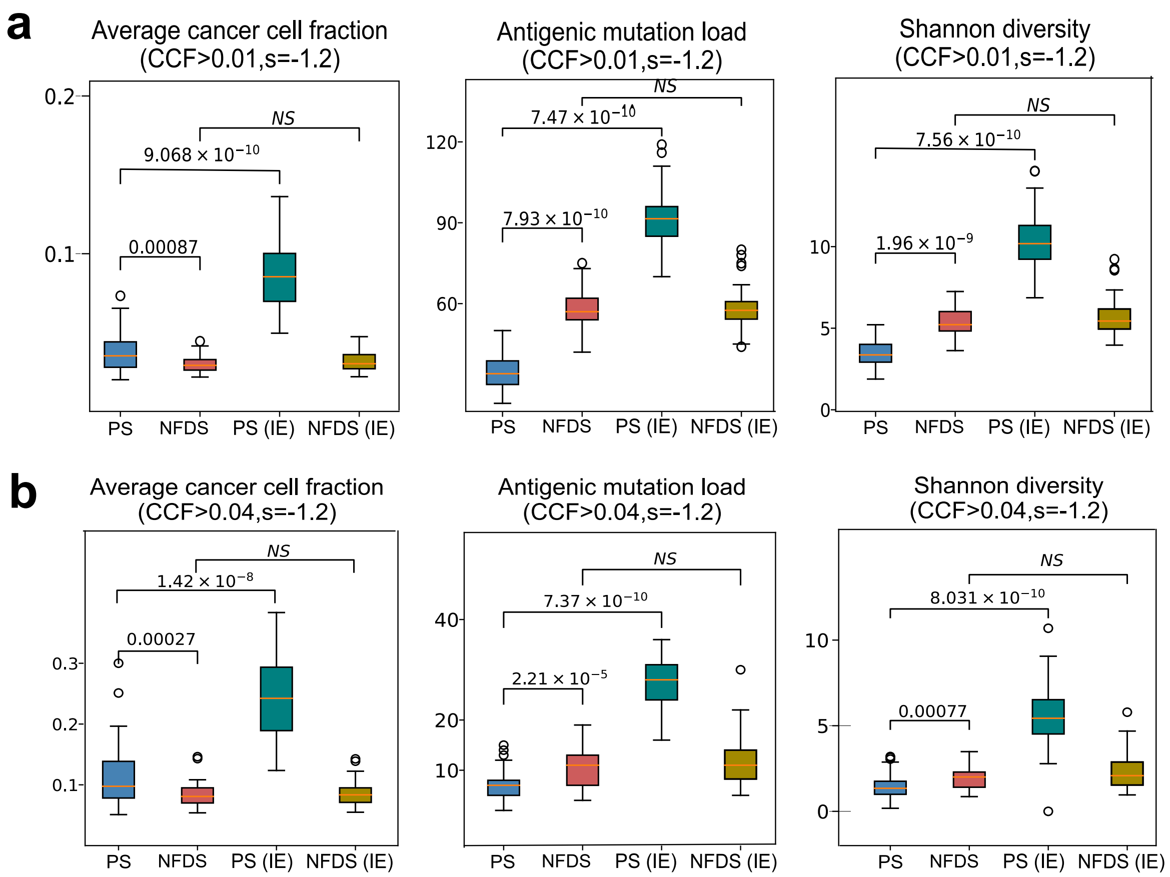


**Supplementary Fig. 8. Cancer cell fraction (CCF), antigenic mutation load and Shannon diversity. a-b,** Box plots showing the antigenic mutations accumulated in 50 simulated tumors with cancer cell fraction (CCF)$> 0.01$ (**a**) and CCF$> 0.04$ (**b**), respectively. The average CCF, mutation load and Shannon diversity of antigenic mutations from 50 simulated tumors under four evolutionary scenarios are shown.

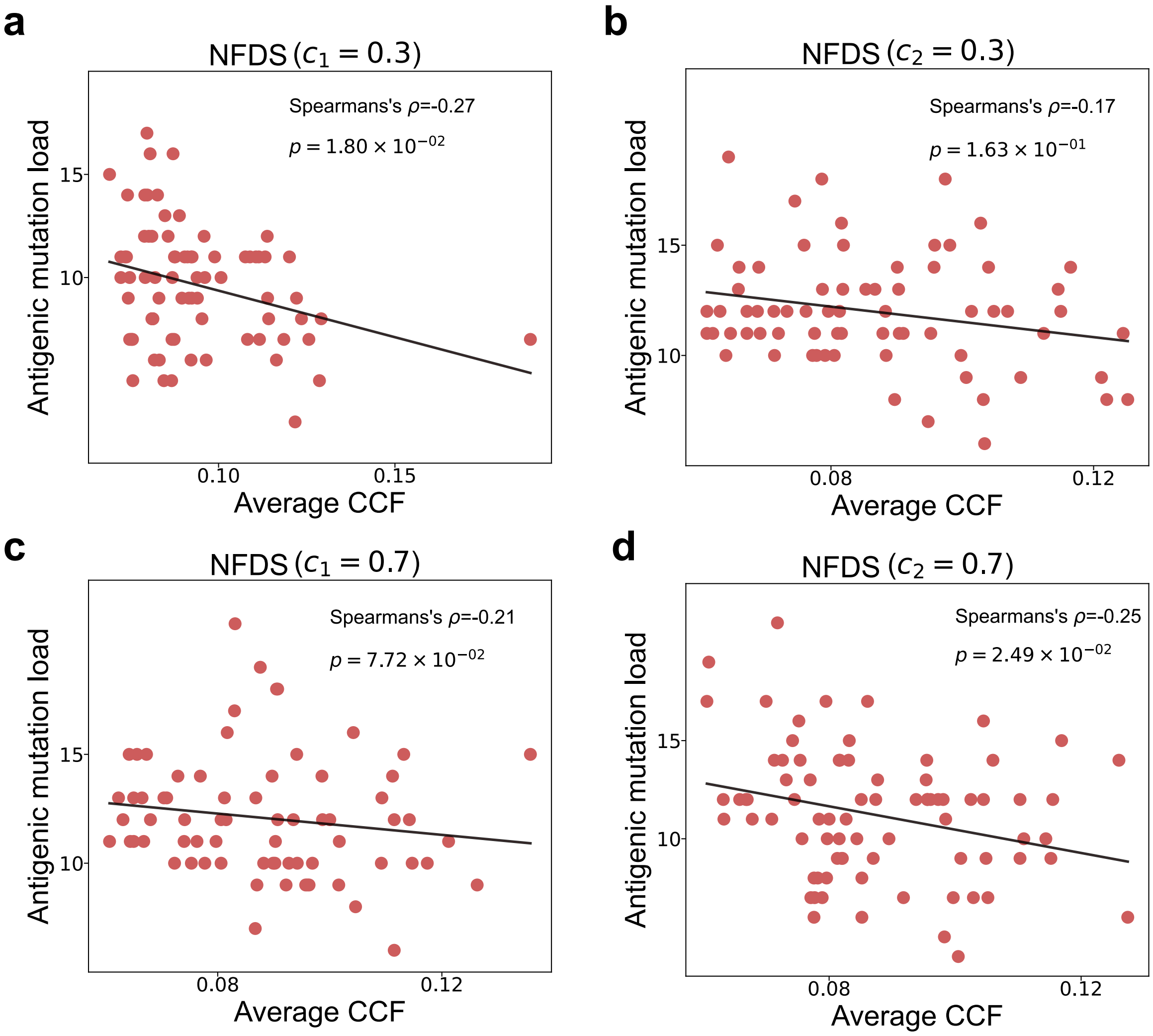


**Supplementary Fig. 9. Correlation analysis between average CCF and subclonal antigenic mutation load. a-b,** Correlation between average CCF and subclonal antigenic mutation load of 100 simulated tumors undergoing NFDS (CCF$> 0.04$) with cell immunogenicity $(c_1)$ equals 0.3 (**a**) and 0.7 (**b**), respectively. **c-d,** Correlation between average CCF and subclonal antigenic mutation load of 100 simulated tumors undergoing NFDS (CCF$> 0.04$) with tumor immunogenicity $(c_2)$ equals 0.3 (**c**) and 0.7 (**d**), respectively.

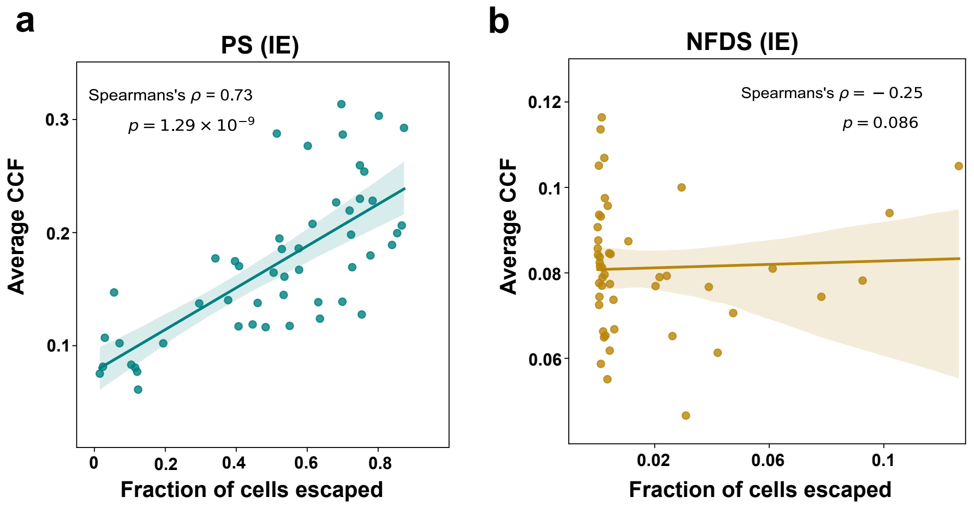


**Supplementary Fig. 10. Correlation analysis between average CCF of neoantigens and fraction of immune-escaped cells. a,** PS (**a**) and NFDS (**b**) with subclonal immune (immune escape probability $p_e = 10^{-4}$, CCF$> 0.04$), respectively.

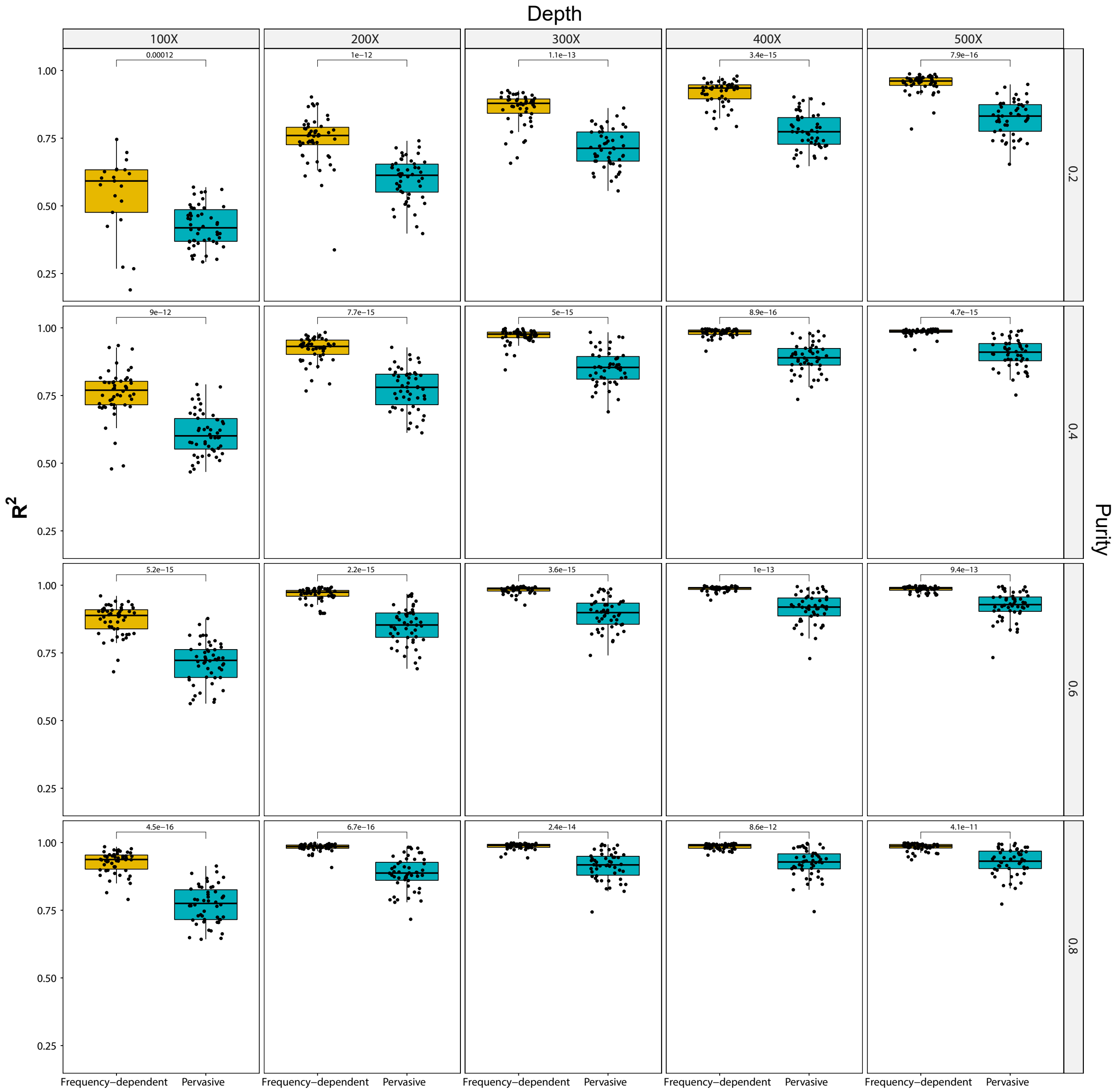


**Supplementary Fig. 11. Identification of neutrality for simulated tumors.** dentification of neutrality with varying sequencing depths and purities in simulations. Simulated depths range from 100x to 500x. Simulated purities range from 0.2 to 0.8. P value, Wilcoxon rank-sum test.

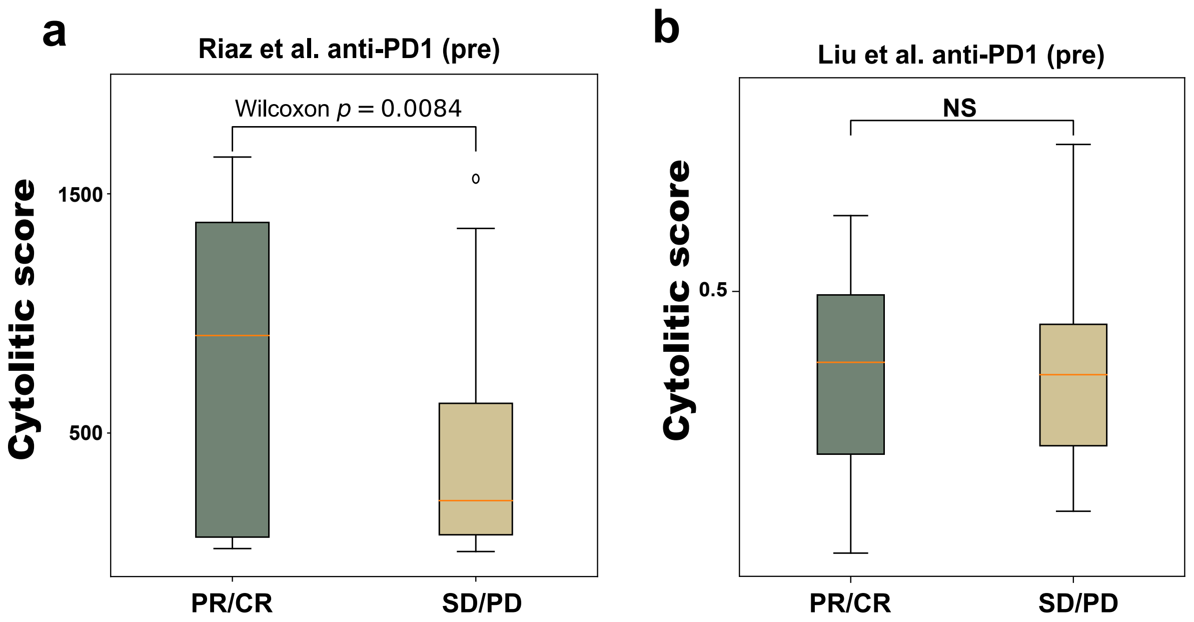


**Supplementary Fig. 12. Immune infiltration in patients. a-b,** Box plots showing the cytolytic score of responders (PR/CR) and non-responders (SD/PD) from Riaz et al. (**a**) and Liu et al.(**b**), respectively.

### 2 Supplementary Note

#### 2.1 The ODE model

To gain deeper understanding of the simulation results, we employed an ordinary differential equation (ODE) model. This model was designed to capture the underlying mechanisms of tumor-immune interactions, drawing inspiration from the classical predator-prey model [1–3]. Since our focus is on hypermutable tumors and neoantigens, we have excluded the transition process of neutral tumor cells to immunogenic cells. To describe the early growth dynamics of an immunogenic tumor lineage $i$, denoted as $I_i$, we can use the following equations:

$$\frac{dI_i}{dt} = bI_i - ksA_{c_i}I_i, \tag{1}$$

where $b$ and $A_{c_i}$ represent the birth rate and antigenicity of the lineage respectively. In order to account for the frequency-dependent mechanism, our model incorporates both negative selection ($s$) and the antigenicity ($A_{c_i}$) of the tumor lineage. We have chosen to describe the exponential growth of tumors in the early stages of tumor progression, without considering size-related effects such as carrying capacity and Michaelis-Menten parameters. This allows us to focus specifically on the initial phases of tumor development. To capture the relationship between the antigenicity of a tumor lineage and the antigenic mutation load, we made the assumption of a linear correlation. Specifically, we observed that higher neoantigen numbers were associated with an increased likelihood of presenting immunogenic cells within the lineage. Additionally, by incorporating the concept of frequency-dependent selection and drawing insights from previous studies [2, 4], we formulated the negative selection ($s$) as $\gamma_i \cdot s_I$ for each immunogenic tumor lineage. Similarly, we represented the antigenicity ($A_{c_i}$) as the sum of all antigenic mutations present in the lineage, denoted as $\sum_k a_k$. Here, $\gamma_i$ denoted the average clonal fraction of antigenic mutations within lineage $i$, while $s_I$ represented the intensity of negative selection.

The dynamics of the immunogenic tumor population can be described by the following equation, which considers the antigenicity of each lineage within the tumor:

$$\sum_i^n \frac{dI_i}{dt} = \sum_i^n I_i \left(b - kA_{c_i}s_I\gamma_i\right), \tag{2}$$

where $I_i$ represents the population of the $i$-th immunogenic lineage of the tumor and $n$ represents the total number of subclones. In growing tumors we have

$$\sum_i^n \frac{dI_i}{dt} = \sum_i^n I_i \left(b - kA_{c_i}s_I\gamma_i\right) > 0. \tag{3}$$

Assuming that each tumor lineage evolves at same mutation rate, we have $I_i \approx I_j$ and set $\gamma_i \approx \gamma_j \approx \gamma$ $(i \neq j)$, where $\gamma$ represents the average CCF of all antigenic mutations in the tumor, which leads to

$$\frac{b}{k \cdot s_I}\frac{n}{\gamma} > \sum_k a_k. \tag{4}$$

The inequality above emphasizes the connection between the antigenic mutation load $\sum_k a_k$ on the right-hand side and $\frac{n}{\gamma}$ on the left-hand side. Drawing inferences from inequality (4), we can conclude that higher levels of intratumoral neoantigen heterogeneity enable a greater capacity for larger neoantigen loads in growing tumors. This implies that tumors with increased intratumoral neoantigen heterogeneity have the potential to accommodate a higher burden of neoantigens as they advance in progression.

We then modeled subclonal immune escape as follows:

$$\begin{aligned} \frac{dI}{dt} &= b \cdot I - ksA_c I - p_e I, \\ \frac{dE}{dt} &= b \cdot E + 2p_e I, \end{aligned} \tag{5}$$

where $I$ represents the population of immunogenic cells, therefore we have $I = \sum I_i$, $A_c = \sum A_{c_i}$. Additionally, we introduce $E$ to represent the population of immune escaped cells, and $p_e$ denotes the probability of immune escape. Based on equation (5), we can infer the following relationship in growing tumors:

$$\frac{dT}{dt} = b \cdot I - ksI \sum_k a_k + b \cdot E + p_e I > 0, \tag{6}$$

where $T$ represents total number of tumor cells. Under the assumption that $T > 0$ and note that $I + E = T$, we have $\frac{I}{T} \leq 1$ and these analyses finally lead to the following inequality:

$$\sum_k a_k < \frac{b + p_e}{ks\frac{I}{T}}. \tag{7}$$

We can deduce from equation (7) that a higher probability of subclonal immune escape allows a tumor to accommodate a larger number of antigenic mutations. Consequently, early subclonal immune escape ensures that the tumor presents a greater number of antigenic mutations with higher clonal fraction (CCF), resulting in an increased antigenic mutation load.

The mathematical analyses presented above highlights different correlation patterns between average CCF and antigenic mutation load in the frequency-dependent selection model and the immune escape model. In tumors subjected to frequency-dependent selection, there exists a negative correlation between the average CCF and antigenic mutation load. However, in tumors exhibiting immune escape mechanisms alone, the average CCF is positively correlated with the antigenic mutation load.
